# Temporal Viral Genome-Protein Interactions Define Distinct Stages of Productive Herpesviral Infection

**DOI:** 10.1101/331272

**Authors:** Jill A. Dembowski, Neal A. DeLuca

**Author notes:** Corresponding author Neal A. DeLuca, Department of Microbiology and Molecular Genetics, University of Pittsburgh School of Medicine, 547 Bridgeside Point II, 450 Technology Dr., Pittsburgh, PA 15219.

## Abstract

Herpesviruses utilize multiple mechanisms to redirect host proteins for use in viral processes and to avoid recognition and repression by the host. To investigate the dynamic interactions between HSV-1 DNA and viral and host proteins, we developed an approach to identify proteins that associate with the infecting viral genome from nuclear entry through packaging. We found that input viral DNA progressed within six hours through four temporal stages where the genomes: 1. interacted with intrinsic and DNA damage response proteins, 2. underwent a robust transcriptional switch mediated largely by ICP4, 3. engaged in replication, repair, and continued transcription, and then 4. transitioned to a more transcriptionally inert state engaging de novo synthesized viral structural components while maintaining interactions with replication proteins. Using a combination of genetic, imaging, and proteomic approaches, we provide a new and temporally compressed view of the HSV-1 life cycle based on genome-proteome dynamics.

## INTRODUCTION

Herpesviruses are a family of highly prevalent eukaryotic viruses that share strong evolutionary relationships with their hosts (McGeoch et al., 2006). They have therefore developed sophisticated mechanisms to invade host cells, alter cellular activities, and redirect host factors for use in viral processes. Knowledge of how herpesviruses manipulate the host to evade intrinsic responses to infection, or to utilize host cell resources to drive productive infection or the establishment of latency, is crucial for understanding their life cycles.

Herpes simplex virus type 1 (HSV-1) is a ubiquitous human pathogen that infects the majority of the human population. Initial productive infection occurs in epithelial cells where the viral genome is prolifically transcribed and replicated, resulting in many new virus progeny. The virus can also gain access to sensory neurons, where it can undergo productive infection or establish reversible latency (Roizman and Whitley, 2013). During latency, most of the viral genome is transcriptionally repressed and no progeny are made. Thus, processes that occur on the viral genome largely determine the outcome of infection.

Nuclear stages of productive infection involve coordinated events occurring on the viral genome, which begin with the transfer of viral DNA into the host nucleus through the nuclear pore shortly after infection (Tognon et al., 1981). Once in the nucleus, the viral genome is subject to the opposing actions of the intrinsic antiviral response mediated at PML nuclear bodies (NBs), and the counteracting functions of viral proteins, particularly ICP0 (Everett et al., 2006; Maul et al., 1993). A coordinated and sequential cascade of expression of three temporal classes of viral genes ensues (Honess and Roizman, 1974, 1975). The transcription of immediate early (IE) viral genes is activated by the viral tegument protein VP16 (Batterson and Roizman, 1983; Campbell et al., 1984), transcription of early and late viral genes is activated by the IE gene product ICP4 (DeLuca et al., 1985; Dixon and Schaffer, 1980; Watson and Clements, 1980), and transcription of late viral genes is coupled to viral DNA replication by an unknown mechanism. IE gene products include regulatory proteins, early gene products include the viral replication machinery, and late gene products mostly comprise the structural components of the virus (Honess and Roizman, 1974, 1975). Replicated DNA is packaged into preassembled capsids, which subsequently exit the nucleus. How these events are staged with respect to the actions of viral and cellular protein complexes acting on the viral genome is unclear. This is mostly due to issues of sensitivity in the measurements of processes occurring during single step growth, and the fact that time of occurrence of crucial events can be obscured by events that are more quantitatively robust.

To examine events that occur on viral genomes, we previously developed an approach based on iPOND (Sirbu et al., 2012) to selectively label replicating viral DNA within infected cells with ethynyl-modified nucleotides (EdC or EdU) to enable the covalent conjugation to biotin-azide or Alexa Fluor-azide (Dembowski and DeLuca, 2015). Biotinylated DNA is purified on streptavidin coated beads followed by the identification of associated proteins by mass spectrometry. Furthermore, Alexa Fluor modified genomes can be imaged in cells relative to specific host or viral proteins. These approaches were used to establish spatiotemporal relationships between specific viral and cellular proteins and the replicating HSV-1 genome and reveal the potential involvement of host factors in processes that occur on nascent viral DNA during relatively late stages of productive infection (Dembowski and DeLuca, 2015; Dembowski et al., 2017). These and other studies also demonstrate that it is possible to track infecting viral genomes that have been pre-labeled with ethynyl-modified nucleotides by imaging approaches (Alandijany et al., 2018; Dembowski and DeLuca, 2015; Sekine et al., 2017; Wang et al., 2013).

Herein we used viral genome purification and imaging approaches to investigate dynamic changes that occur on the original infecting viral genome during distinct stages of infection. HSV-1 structural and tegument proteins from the infecting virus are associated with the infecting viral genome early during infection. As infection proceeds, the same structural proteins are synthesized in the infected cell and then again become associated with viral genomes. Therefore, we utilized stable isotope labeling of amino acids in cell culture (SILAC) to differentiate between proteins that originate in the infecting virion and proteins that were synthesized within the infected cell. Additionally, we performed this analysis on both wild type virus and a virus that does not synthesize ICP4, to investigate changes that mediate a robust transcriptional switch early during infection. Using a combination of genetic, imaging, and proteomic approaches, we have tracked the fate of input viral genomes within the nuclei of host cells throughout the entire course of infection and defined several crucial steps that occur early in the productive HSV-1 life cycle.

## RESULTS

### Input Viral Genomes Can Be Tracked from Nuclear Entry Through Packaging

To investigate the protein landscape associated with input viral DNA, wild type HSV-1 (KOS) stocks were prepared in the presence of EdC to label viral genomes, which enables subsequent imaging or purification of input viral DNA after infection. EdC-labeling resulted in an approximately three-fold increase in the genome/plaque forming unit (PFU) ratio (Table S1), suggesting that incorporation of EdC into viral DNA results in a modest decrease in infectivity.

To demonstrate the sensitivity and specificity of viral genome labeling, Vero cells were infected with EdC-labeled KOS (KOS-EdC) and fixed at various times after infection. Fixed cells were subject to click chemistry and immunofluorescence to visualize the relative location of input viral DNA and ICP4 within the host nuclei (Figure 1A). At 1 hour post infection (hpi), viral genomes were observed at the perimeter of the nuclear membrane as they entered into the nucleus through the nuclear pore. By 2 hpi, ICP4 was expressed and colocalized with most if not all viral genome foci. From 3-12 hpi, after the onset of viral DNA replication, ICP4 foci representing replication compartments grew in size, while input genomes contained within these foci could still be distinguished as discrete puncta. At later times, input viral DNA appeared to coalesce and migrate to the perimeter of replication compartments. Together these data demonstrate the specificity of the click chemistry approach for tracing the fate of the input viral genome throughout the course of infection (Figure 1B).

**Figure 1.**
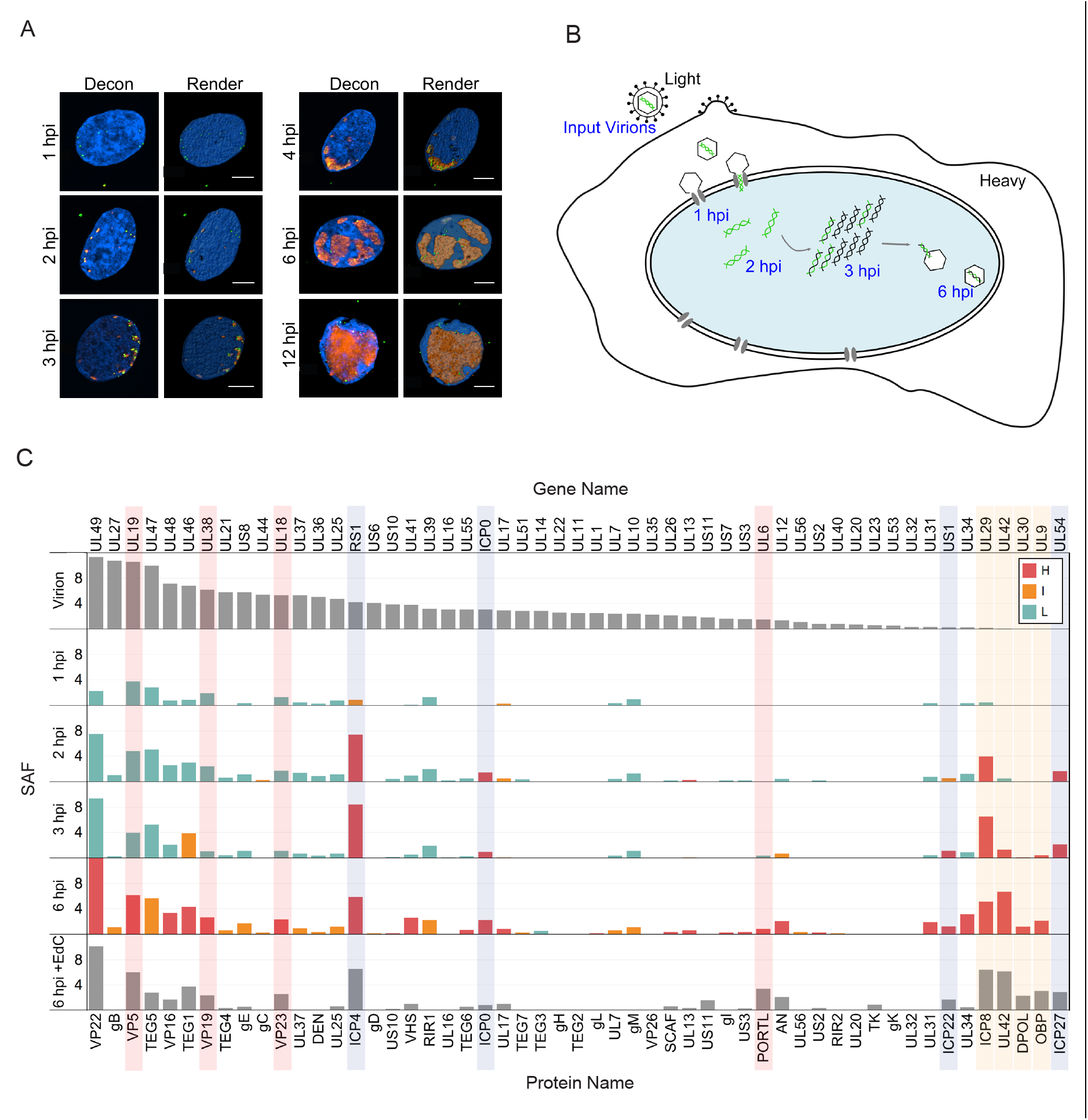
The Fates of Input Viral Genomes can be Tracked within Infected Cell Nuclei. A. Visualization of input viral DNA. Cells were fixed at indicated times after infection with KOS-EdC (1-12 hpi). Viral genomes (green) and ICP4 (red) were imaged relative to host nuclei (blue). Optical sections were deconvolved (Decon) to generate high resolution images and rendered (Render) to illustrate the positions of input viral DNA. Scale bars represent 5 μm. B. Model depicting the fate of input viral genomes during distinct stages of infection. Input viral DNA is shown in green and replicated viral DNA in black. For MS experiments, input virions and the infected cell contain proteins labeled with light or heavy amino acids, respectively. C. Abundance of viral genome associated viral proteins throughout infection. Viral genome associated viral proteins were detected by MS after a 1, 2, 3, or 6 hour infection with KOS-EdC. SAFs were plotted relative to individual proteins associated with mature virions (Virion) and viral replication compartments that were labeled with EdC from 4-6 hpi (6hpi + EdC) (Dembowski et al., 2017). Proteins were distinguished as either heavy (H, red), light (L, teal), or intermediate (I, orange) by SILAC analysis. In cases were SILAC analysis was not carried out, bar graphs are shown in gray. IE viral proteins are highlighted in purple, viral replication proteins in tan, and capsid proteins in red. Data represent results from one of two biological replicates. See also Tables S1 and S2, Figures S1 and S2.

To investigate the ordered protein interactions that occur on input viral DNA, human MRC-5 fibroblast cell nuclei were harvested at specific times after infection with KOS-EdC and viral DNA was covalently attached to biotin and purified using streptavidin-coated beads followed by mass spectrometry (MS) to identify associated proteins (Figure 1C, Table S2). To compare the relative abundance of identified proteins, spectral abundance factors (SAF: spectral counts/molecular weight) were calculated for each protein and plotted in order of relative abundance in mature virions (Figure 1C, Virion). For comparison, proteins found to associate with viral replication compartments were also graphed (6 hpi + EdC) (Dembowski and Deluca, 2017). SILAC was carried out to distinguish between factors that originated in the infecting virion (light amino acids) and factors that were expressed within the infected cells either prior to or during infection (heavy amino acids). Relative intensities of amino acids were compared (Figure 1C) and input genome associated proteins were distinguished as heavy (H), light (L), or intermediate (I) and to have therefore originated in the infected cell, virion, or both. SILAC MS analysis of viral proteins was highly reproducible (Figure S1A).

At 1 hpi, viral genomes associated with light capsid and tegument proteins that were brought into the cell with the infecting virion. Immediate early viral gene products (ICPs 0, 4, 22, and 27) are expressed by 1 hpi (Harkness et al., 2014) and were found to associate with infecting viral genomes by 2 hpi (Figure 1C, highlighted in purple). ICP8 was the first viral replication factor to associate by 2 hpi and additional replication factors were detected by 3 hpi (UL42, UL30, UL9, highlighted in tan), the time at which viral genomes begin to replicate (Dembowski et al., 2017). Replication factors are not abundant in the virion, and were therefore expressed de novo subsequent to infection and generally contained heavy amino acids. At later times (6 hpi), viral genomes were found to associate with newly expressed viral structural proteins, including capsid proteins VP5, VP19, VP23, and UL6 (highlighted in red). A clear transition from light to heavy capsid proteins was observed by 6 hpi, and nascent capsid proteins associated with these genomes in roughly the same relative abundance as they constitute intact capsids (Figure S1B) (Gibson and Roizman, 1972; Newcomb et al., 1993). Therefore, some population of the input viral DNA was repackaged by this time.

Co-staining of input genomes, ICP4, and the major viral tegument protein (VP5) demonstrates that labeled input genomes are released from capsids docked at the nuclear membrane at early stages of infection (1 hpi); these genomes associate with ICP4 by 2 hpi; nascent VP5, a late gene product, accumulates in the nucleus by 4 hpi; and input genomes colocalize with VP5 associated with replication compartments by 6 hpi (Figure S2). Taken together, temporal viral genome-viral protein interactions observed in these studies are consistent with known events in the virus life cycle and demonstrate the sensitivity, specificity, and reproducibility of the input viral genome purification approach.

### Host Proteins Associated with Input Genomes upon Nuclear Entry

Host proteins in the MS datasets of affinity purified input genomes are listed in Table S2. To further demonstrate the reproducibility of this assay to determine the relative abundance of factors associated with viral DNA, the SAF values of individual proteins from duplicate experiments were plotted to determine the Pearson correlation coefficient of replicate experiments (Figure S3). In all cases, the correlation coefficient was at least 0.92, demonstrating a linear relationship between data points and a general consistent trend in relative yield of individual factors using this approach. Although many experimental variables govern whether a protein will be captured and identified using this technique, results are consistent between replicate experiments and there is high confidence in the presence of identified factors.

No host factors were reproducibly found to contain peptides labeled with light amino acids. Therefore, we conclude that cellular proteins associated with viral genomes at early stages of infection do not originate from the infecting virus particle. Potential interactions amongst viral genome associated host factors identified by MS were illustrated using the STRING protein-protein interaction network database (Snel et al., 2000). One hour after infection, host proteins identified to associate with viral DNA include the catalytic subunit of host Pol II (POLR2A), factors that play roles in transcription regulation and RNA processing (INTS1, USP39, SRRT, DDX23, THOC7), core components of PML NBs (PML, SP100, SUMO2), factors involved in the regulation of chromatin structure (HP1BP3, HIST1H1A, HIST1H1E, CHD4, CSNK2A1, SUPT16H, SMARCC2, SMARCA4, TRRAP), and factors that are recruited to damaged DNA (PARP1, PARP14, RPA1, LIG3). Identified host factors illustrate the processes that occur on HSV-1 genomes shortly after entry into the nucleus: 1) transcription of IE viral genes, 2) association with PML NBs and components of cellular chromatin, and 3) recognition by the host cell as DNA damage.

To determine if multiple processes occur on each genome, or if these results reflect the existence of mixed populations of viral DNA engaged in different processes at 1 hpi, we carried out co-staining for a protein involved in viral repression (PML) and the core subunit of Pol II (POLR2A) (Figure 2B). We demonstrate that individual viral genome foci are associated with both PML and Pol II, but that PML and Pol II do not colocalize with each other. This is consistent with the observation that viral genomes are juxtaposed to PML NBs at this time (Ishov and Maul, 1996). We cannot distinguish between whether individual foci contain more than one genome. However, because these foci appear to originate from a single capsid focus (Figure S2), we hypothesize that the viral genome foci represent individual genomes at early times post infection (1-2 hpi). Taken together, viral genomes are recognized by the cell as DNA damage early during infection and are associated with PML NBs. However, portions of the viral genome can escape repression to enable the transcription of IE viral genes.

**Figure 2.**
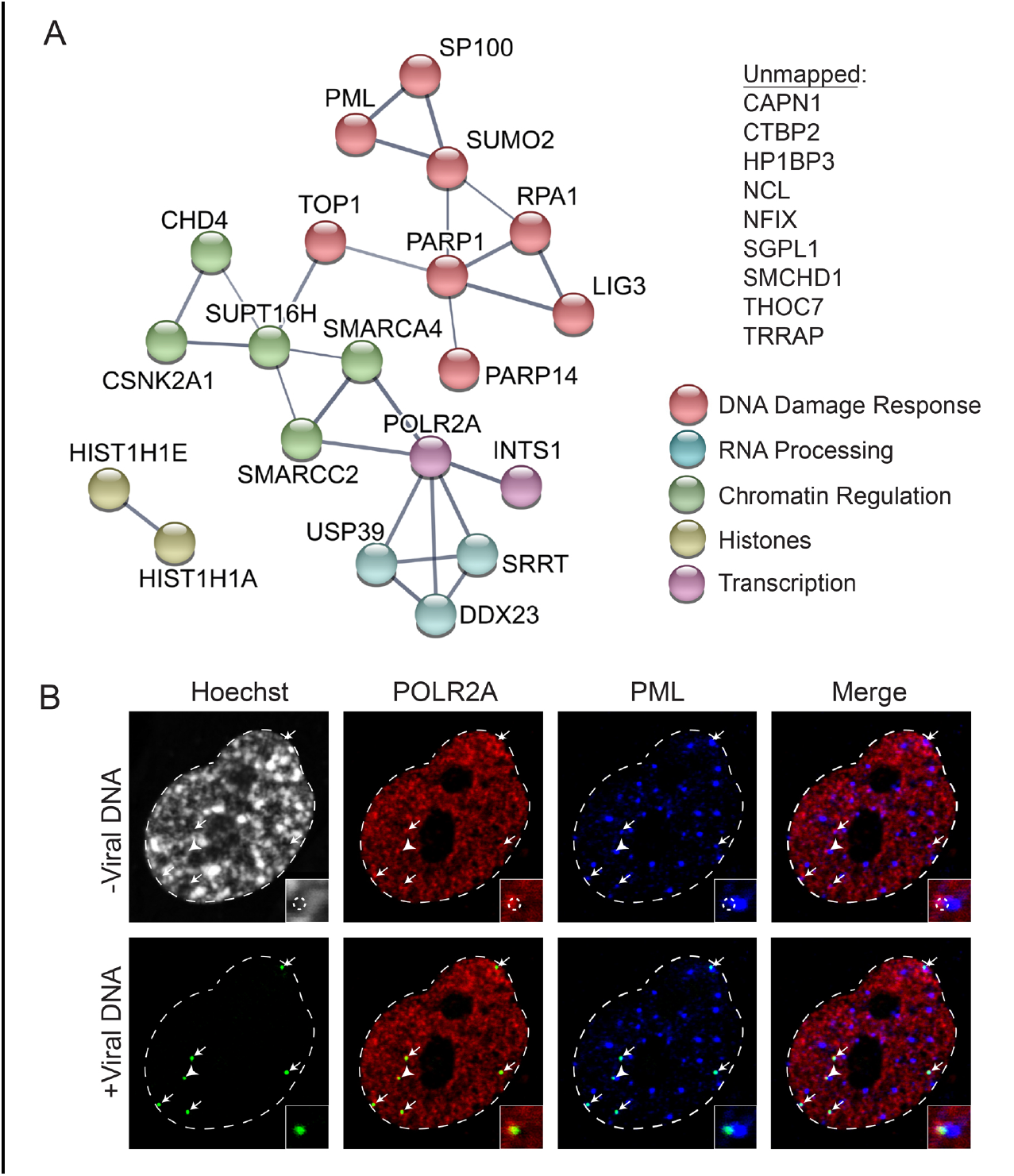
Host DNA Damage Response Factors and Pol II Associate with Input Viral Genomes by 1 hpi. A. Graphic illustration of predicted physical and functional interactions between human proteins that associate with HSV-1 genomes at 1 hpi with KOS-EdC. Colors indicate the biological processes in which identified proteins are likely involved and the unmapped list includes proteins that were not mapped using STRING. B. Input viral genome foci simultaneously associate with Pol II and PML. KOS-EdC infected MRC-5 cells were fixed at 1 hpi and subject to click chemistry to label input viral DNA (green), Hoechst staining to label nuclei, and indirect immunofluorescence to label Pol II (POLR2A, red) and PML (blue). All panels represent different views of the same nucleus, which is outlined in white. Arrows indicate the location of input viral genomes and the corner box includes a 3x zoomed in image of the area indicated by the arrow head. Viral DNA was omitted from the top panel (-Viral DNA) and the location of input viral genome foci is outlined by a dashed circle in the zoomed in image. Viral DNA foci were included in the bottom panel (+ Viral DNA). See also Figure S3 and Table S2.

### Robust Transcription Factor Recruitment to Viral Genomes Occurs Coincident with the Binding of ICP4

After two hours, PML NBs are dispersed through the actions of ICP0, and ICP4 associates with the viral genome to activate transcription of early viral genes. In this study, nascent IPC4 was found to associate with input viral genomes (Figure 1C), and PML components (PML, SP100, SUMO2) were no longer detected by 2 hpi (Figure 3A). At this time, several host factors involved in host cell transcription were found to associate with viral DNA. These include components of the host Mediator (MED1, 6, 12, 14, 16, 17, 20, 23, 24, 25, 27) and Integrator (INTS1, 2, 3, 4, 6, 7,10 and CPSF3L) complexes, as well as factors that regulate transcription elongation (SSRP1, SPT16H, SUPT5H, SUPT6H) and RNA processing. With the exception of POLR2A, INTS1, and THOC7, associated transcription and RNA processing factors were not detected before 2 hours and were therefore potentially recruited through the actions of ICP4, another IE viral gene product, or as a result of alterations in viral genome architecture. It is also possible that small amounts of these complexes are recruited to the genome earlier through the action of VP16, but are present below the limits of detection. We previously demonstrated that ICP4 interacts with the Mediator complex and was required for the recruitment of Mediator components to viral promoters (Lester and DeLuca, 2011).

**Figure 3.**
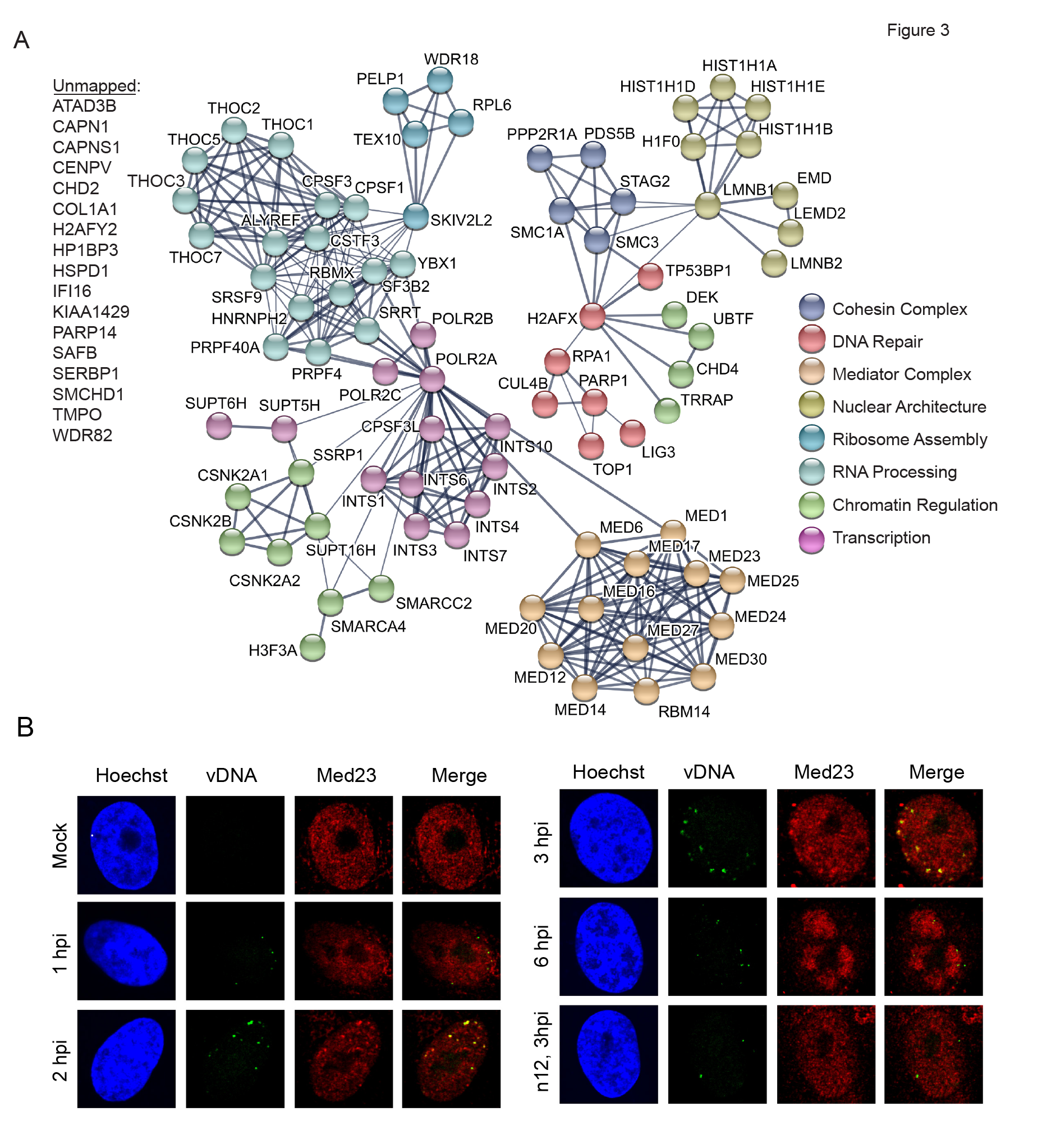
Robust Recruitment of Host Transcription Factors to Input Viral DNA. A. Illustration of predicted physical and functional interactions between human proteins that associate with HSV-1 genomes at 2 hpi with KOS-EdC. B. Input viral genome foci associate with Med23 starting around 2 hpi. KOS-EdC or n12-EdC infected MRC-5 cells were fixed at the indicated times and subject to click chemistry to label input viral DNA (green), Hoechst staining to label nuclei (blue), and indirect immunofluorescence to label Med23 (red). See also Figure S3 and Table S2.

Furthermore, the IE gene product ICP22 is required for the recruitment of SSRP1 to viral DNA (Fox et al., 2017). To verify the timing of recruitment of Mediator to viral DNA, we co-stained infected cells for infecting viral genomes and the Mediator component Med23 (Figure 3B). Med23 did not colocalize with viral genomes by 1 hpi but did colocalize by 2 hpi, validating the MS results.

Furthermore, by 2 hpi, TP53BP1 and IFI16 were also found to associate with viral genomes, as well as components of the nuclear lamina (LEMD2, EMD, LMNB1, LMNB2) and the cohesin complex (SMC1A, SMC3, STAG2, PDS5B). Interestingly, the nuclear lamina may play a role in the reduction of heterochromatin on viral genes (Silva et al., 2008). Taken together, there is an obvious switch in viral genome architecture that occurs between 1 and 2 hpi that likely mediates the onset of early viral gene expression and sets the stage for viral genome replication.

### ICP4 Facilitates Transcription Factor Recruitment

To investigate the role ICP4 plays in the recruitment of host transcription factors to viral DNA, stocks of the ICP4 mutant, n12 (DeLuca and Schaffer, 1988), were prepared by propagating the virus in the presence of EdC. EdC labeling of n12 in the ICP4 completmenting cell line E5 resulted in a two-fold increase in the genome/PFU ratio (Table S1). Therefore, as observed for wild type KOS, viral genome labeling resulted in a modest decrease in n12 infectivity. To verify that EdC-labeling was specific, n12-EdC infected cells were subject to immunofluorescence (Figure S4). Viral genomes were observed at the perimeter of the nucleus at 3 hpi (n12, Vero, 3 hpi) and did not progress to form replication compartments unless ICP4 was supplied in trans (n12, E5, 6 hpi). Therefore, the analysis of n12-EdC infection should enable the investigation of changes that occur on the viral genome as a consequence of ICP4 association.

MRC-5 cells were infected with n12-EdC and proteins that associated with the genome at 3 hpi were determined as described above. Identified viral proteins were graphed relative to viral proteins found to associate with KOS-EdC viral genomes at this time (Figure 4). A small amount of ICP4 was purified with n12 viral genomes. We conclude that this population of ICP4 was carried into the cell as part of the viral tegument because it was enriched in light amino acids. MS of purified virions also revealed that ICP4 is a component of mature n12 virions, which contain the same protein composition as KOS virions (Figure S5). n12 infected cells do not efficiently express early viral genes including, ICP8, UL42, UL9, or UL30 and as a consequence these proteins were not abundantly associated with n12 viral genomes which do not undergo viral DNA replication in noncomplementing cells (DeLuca and Schaffer, 1988).

**Figure 4.**
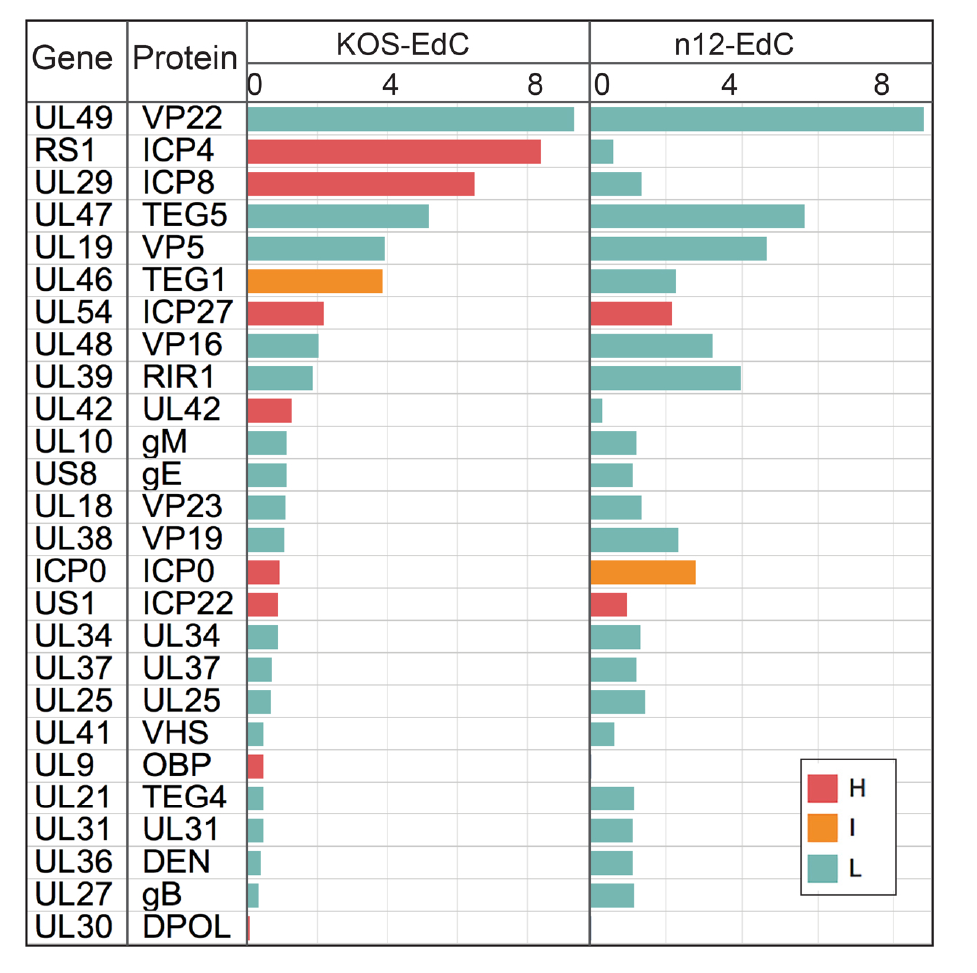
ICP4 is Required for the Expression and Subsequent Association of Early, but not IE, Viral Gene Products with Input Viral Genomes. Viral genome associated viral proteins were detected by MS at 3 hpi with KOS-EdC or n12-EdC. Proteins were distinguished as either heavy (H,red), light (L, teal), or intermediate (I, orange) by SILAC analysis. See also Tables S1 and S2, Figures S1, S4, and S5.

To establish the requirement of ICP4 for the recruitment of host factors to viral DNA, we compared the average SAFs of human proteins enriched on n12 and KOS viral genomes at 3hpi (Figure 5A). Proteins that fell within the 90% confidence interval of the linear regression line were considered to be enriched on both KOS and n12 viral genomes. Proteins that fell outside of this confidence interval were considered to be enriched on KOS and not n12 viral genomes (red) or enriched on n12 and not KOS viral genomes (green). The identified proteins were further grouped based on their biological function and the average SAF values of proteins within each group were compared between KOS and n12 infected cells (Figure 5B). Processes that were enriched on KOS over n12 genomes are shown in red and processes that were enriched on n12 over KOS genomes are shown in green. To further illustrate the differences in individual proteins that were more enriched on KOS or n12 viral genomes (Figure 5A), STRING maps were generated (Figures 5C, D). From these data, we conclude that ICP4 is required for the recruitment of the Mediator complex (Figure 5C, dark red) to viral DNA, as well as several transcription elongation factors (tan: CDK9, SUPT5H, SUPT6H) and factors that are enriched at viral replication forks (dark orange: TOP2A, PCNA, MRE11) (Dembowski et al., 2017). However, in the absence of ICP4 there is an increase in factors that recognize DNA damage (Figure 5D, light green: PARP9, DTX3L, PARP1, XRCC6, RPA1, RPA2), RNA processing factors (light blue), the FACT complex (dark blue: SSRP1, SPT16H), as well as histones and chromatin remodeling factors (dark green). Consistent with MS results, we did not observe the recruitment of Med23 to n12 viral genomes by immunofluorescence (Figure 3B). Taken together, ICP4 association with viral DNA triggers a significant change in viral genome architecture resulting in a transition from a state involving chromatin repression and recognition as DNA damage to a state associated with robust transcription and viral DNA replication.

**Figure 5.**
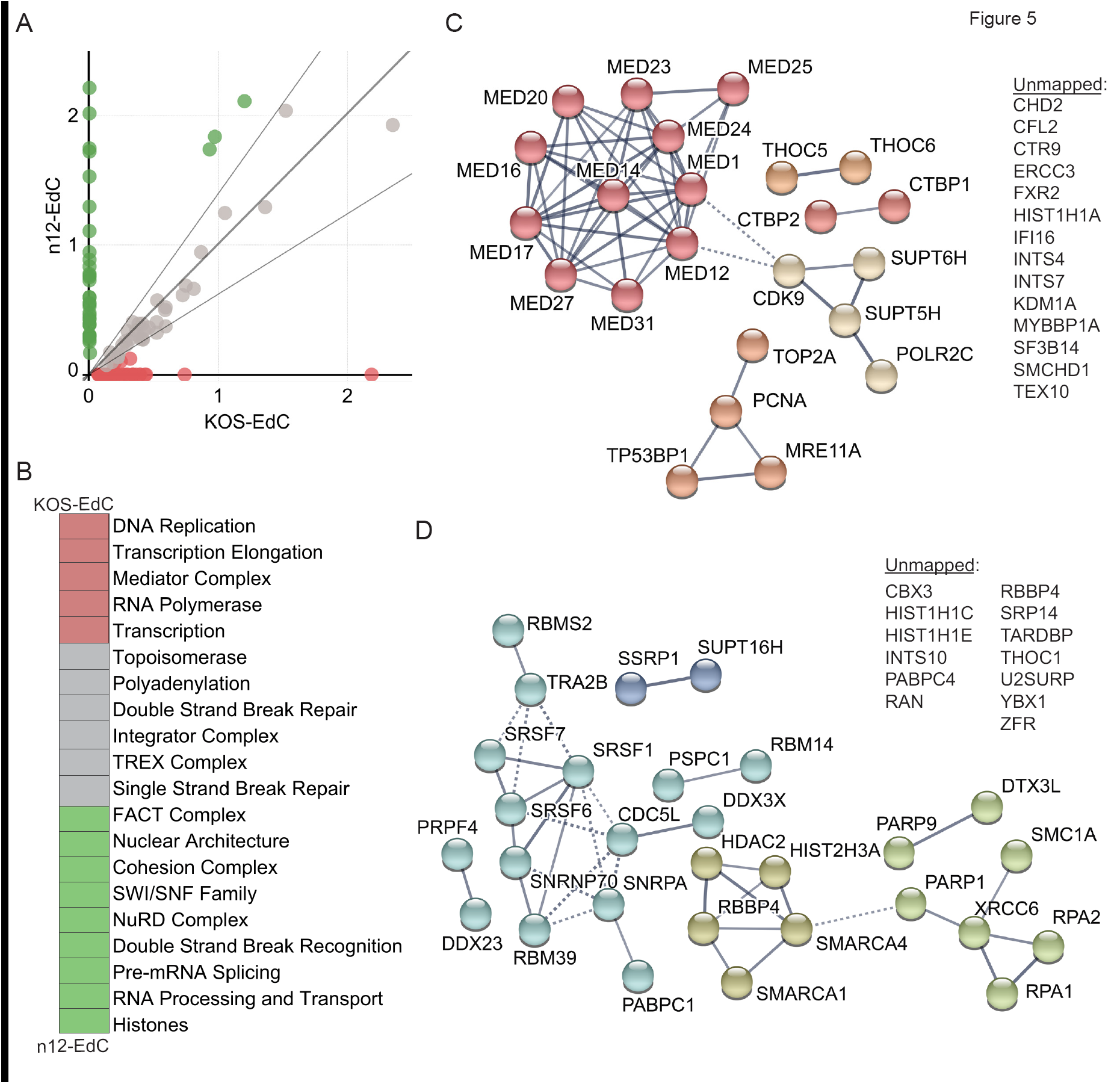
ICP4 is Required for the Recruitment of Host Transcription and Replication Factors to Input Viral DNA. A. Relative enrichment of human proteins associated with n12-EdC versus KOS-EdC genomes. Each point represents an individual viral genome associated host protein with the average spectral abundance on KOS-EdC at 3 hpi plotted on the x-axis and the average spectral abundance on n12-EdC at 3 hpi plotted on the y-axis. The linear regression line was forced through zero and the 90% confidence interval is shown. Proteins that fell outside of this confidence interval were considered to be relatively more enriched on KOS (red) or n12 (green) viral genomes. B. Viral genome associated host proteins were grouped based on their biological function and the average SAF values of proteins within each group were compared between KOS-EdC and n12-EdC infected cells. Processes that were >1.5 fold more enriched on KOS viral genomes are shown in red and processes that were >1.5 fold more enriched on n12 viral genomes are shown in green. C-D. Graphic illustration of predicted physical and functional interactions between human proteins that were relatively more enriched on KOS-EdC viral genomes compared to n12-EdC (C) or n12-EdC viral genomes compared to KOS-EdC (D). See also Table S2 and Figure S3.

### Host Proteins Associated with Input Viral Genomes at the Onset of Viral DNA Replication

We previously detected viral DNA replication in infected MRC-5 cells as early as 3 hpi (Dembowski et al., 2017). Furthermore, this was the earliest time at which viral replication factors UL30, UL9, and UL42 were detected to associate with viral DNA (Figure 1C and Table S2). To investigate host factors that associate with infecting viral genomes immediately after or during the onset of viral DNA replication, host proteins associated with input viral DNA at 3 hpi were identified. At this time, we observed the association of the TFIIH component ERCC3, the Pol II kinase CDK9, and Mediator component Med31 with viral DNA (Figure 6A). ERCC3 and CDK9 were previously shown to associate with replicated HSV-1 DNA (Dembowski et al., 2017), are known to have roles in the active transcription of cellular genes, and may therefore play a role in activating late viral gene expression. ERCC3, CDK9, and Med31 were not found to associate with viral DNA, at least within the limits of detection by this method, in the absence of ICP4 (Figure 5C), suggesting that ICP4 may play a role in recruiting these factors.

**Figure 6.**
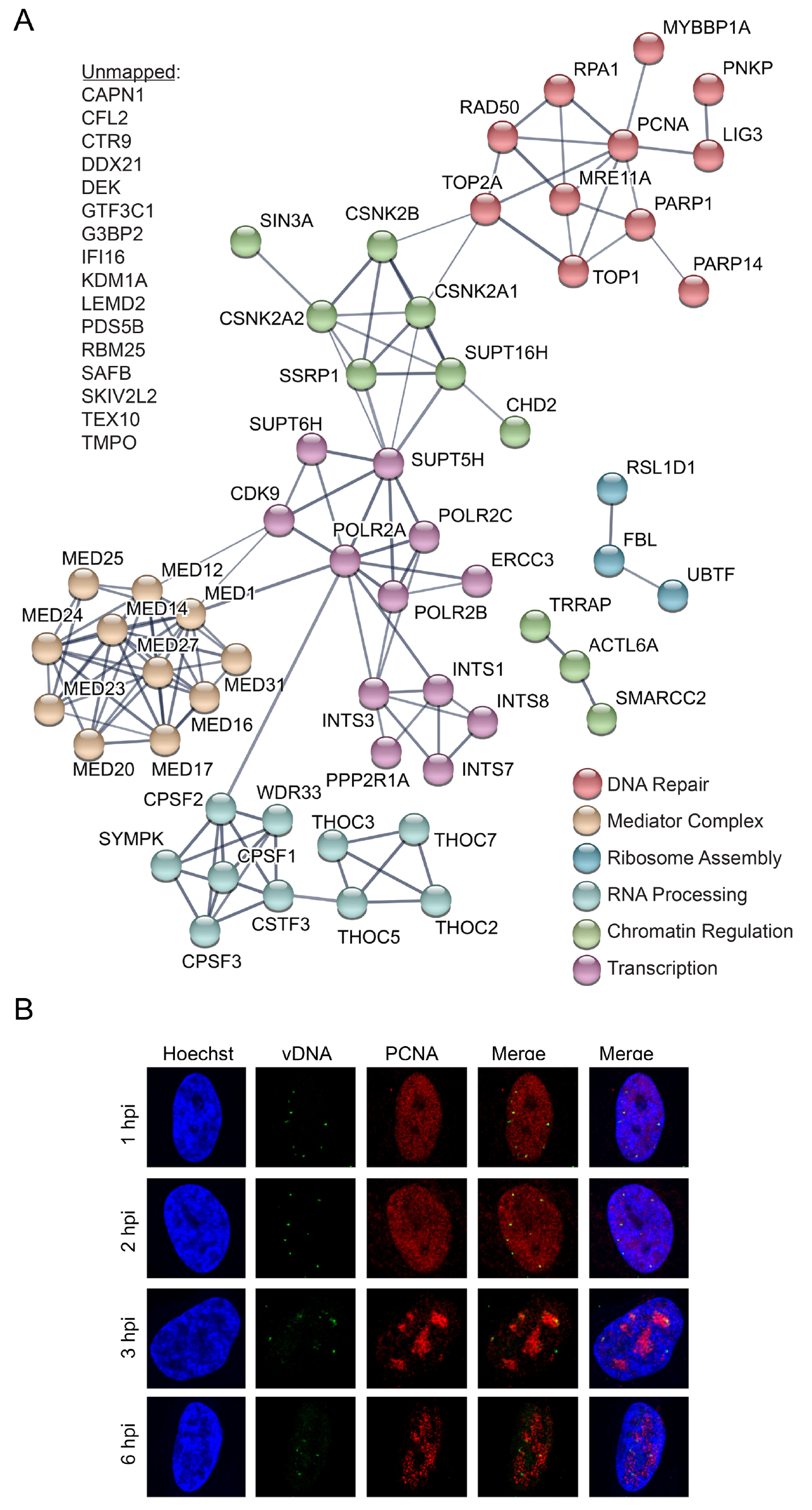
Additional Host Factors are Recruited to Input Viral DNA at 3 hpi. A. Graphic illustration of predicted physical and functional interactions between human proteins that associate with KOS-EdC genomes at 3 hpi. B. Input viral genome foci associate with PCNA at 3 hpi. KOS-EdC infected MRC-5 cells were fixed at 1, 2, 3, or 6 hpi and subject to click chemistry to label input viral DNA (green), Hoechst staining to label nuclei (blue), and indirect immunofluorescence to label PCNA (red). See also Table S2 and Figure S3.

After 3 hours, we also observed the association of PCNA and the topoisomerase subunit TOP2A with viral DNA (Figure 6A and B). The MRN double strand break repair complex members MRE11A and RAD50 also associated at this time (Figure 6A). Consistent with previous observations, all of these factors have been shown to associate with replicated viral DNA (Dembowski and DeLuca, 2015; Dembowski et al., 2017). The functions of host repair proteins and PCNA on replicating HSV-1 DNA are not known, however it has previously been demonstrated that PCNA and MRE11 are required for efficient viral DNA replication (Lilley et al., 2005; Sanders et al., 2015). Taken together, at the onset of viral DNA replication, another unique set of factors associate with input viral genomes. These factors likely play a role in replication-coupled processes such as the repair of damaged DNA, recombination, or activation of late gene transcription.

### Heterogeneity of Genomes at Late Times Post Infection

At 6 hpi, we observed the robust association of infecting viral genomes with many host factors (Figure 7A, Table S2). These include the cohesin complex, cytoskeletal proteins, components of the nuclear lamina, DNA repair proteins, RNA processing factors, and factors that regulate chromatin structure. Newly associated proteins include recombination (RECQL), base excision repair (BER) (APEX, XRCC1), and mismatch repair (MMR) (MSH2) proteins, suggesting that viral genomes undergo repair and recombination at this time. Input viral genomes continue to associate with viral replication factors at 6 hpi and therefore at least some population of input viral DNA continues to undergo DNA replication. It is likely that BER and MMR occur on nascent viral DNA associated with input viral genomes in the act of DNA replication. Consistent with this hypothesis, we previously observed the association of these factors with replicated viral DNA (Dembowski and DeLuca, 2015). Another population of viral genomes appear to be packaged into capsids composed of nascent viral proteins (Figure 1C) and it may also be this population that associates with microtubule associated proteins (Figure 7A MAP1A, MAP1B, DYNLL1, CKAP5) at this time. Microtubule associated proteins may facilitate the transport of nascent nucleocapsids. One striking observation is that input genomes exhibit reduced association with several transcription factors including the Mediator complex, Integrator complex, and Pol II but increased or continued association with factors that regulate chromatin architecture, including NuRD, B-WICH, Swi/Snf, and FACT (Figure 7B). While the functions of chromatin remodeling factors on viral DNA at this time are unknown, the decrease in Pol II levels suggests that transcription is likely reduced from input viral genomes. Taken together, input viral genomes are present in mixed populations by 6 hpi, whereby some genomes continue to replicate, while others are processed and packaged into virions.

**Figure 7.**
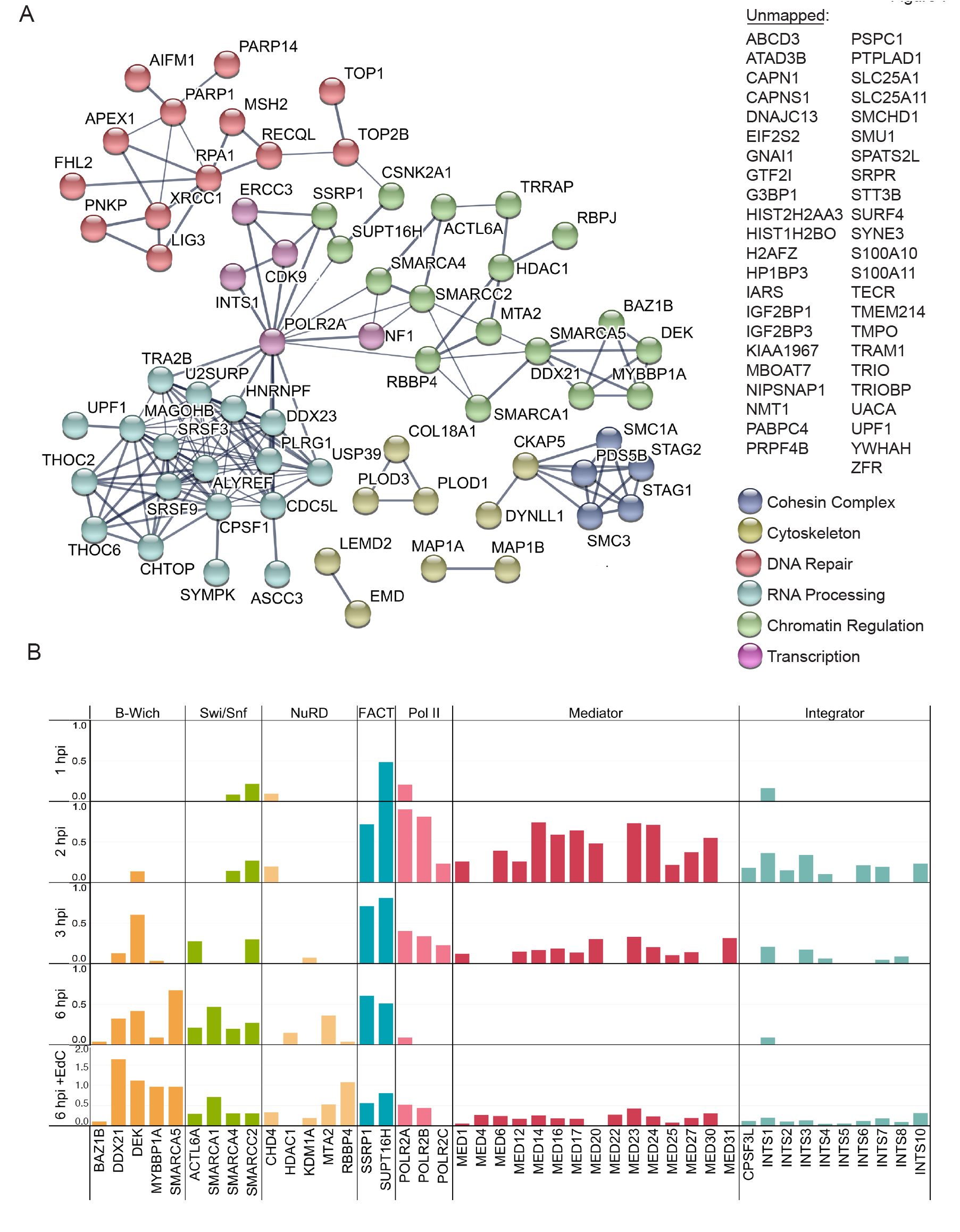
Levels of Host Transcription Factors are Selectively Reduced on Input Viral DNA at 6 hpi. A. Graphic illustration of predicted physical and functional interactions between human proteins that associate with HSV-1 genomes at 6 hpi with KOS-EdC. B. Spectral abundance (SAF) of select viral genome associated host proteins that function in transcription regulation. Viral genome associated host proteins were detected by MS at 1, 2, 3, or 6 hpi with KOS-EdC. Proteins that associate with viral replication compartments that were labeled with EdC from 4-6 hpi (6hpi + EdC) are also shown (Dembowski et al., 2017). Note that KDMA1 is associated with the NuRD complex but is not a core component. See also Table S2 and Figure S3.

## DISCUSSION

Infecting viral genomes are acted on by host and viral proteins to facilitate sequential steps in infection. Here, we developed and utilized an approach to investigate the origin and temporal association of viral and host proteins with input HSV-1 genomes from nuclear entry through repackaging into nascent capsids within 6 hours (Figure 1B). These studies provide a new and temporally compressed view of the life cycle of HSV-1 based on the dynamics of the genomic proteome, and provide new evidence for the involvement of specific host factors in each step.

### Tracking Viral Protein Dynamics

Early during infection, viral genomes associate with capsid and tegument proteins that originate from the infecting virus (Figure 1C, 1-3 hpi, capsid proteins highlighted in red). However, compared to purified virions (Virion), viral genomes purified after infection are not associated with an abundance of viral glycoproteins and therefore enveloped virus particles. It has previously been demonstrated that the click reaction can only access viral genomes after release from the capsid through the nuclear pore (Sekine et al., 2017) or partial denaturation to disrupt capsid integrity (Alandijany et al., 2018). Therefore, at early times, viral genome associated capsid proteins isolated from nuclei likely associate with viral DNA docked and uncoating at the nuclear pore (Figure S2, 1-3 hpi). Later during infection, nascent capsid proteins that were synthesized after infection and therefore contain heavy amino acids associate with viral genomes (Figure 1C, 6 hpi) in a similar relative abundance as they constitute intact capsids (Figure S1B). Therefore, while some of input genome foci present at 6 hours may represent genomes that have not progressed through the infection process as previously proposed (Sekine et al., 2017), our data support a model whereby some population of input viral DNA begins to be repackaged by 6 hpi, putting an upper limit on the minimum time necessary for completion of the nuclear events in productive infection.

Identified interactions with regulatory viral factors are consistent with previous information regarding the virus life cycle. VP16 from the infecting virion associates with viral DNA early during infection (Figure 1C) to mediate expression of IE viral genes (Batterson and Roizman, 1983; Campbell et al., 1984). Nascent IE gene products associate with viral genomes as early as 2 hpi (Figure 1C, IE proteins highlighted in purple). IE gene products, including ICP4, drive the expression of early genes (DeLuca et al., 1985; Dixon and Schaffer, 1980; Watson and Clements, 1980). Early genes encode the viral replication machinery (highlighted in tan), which are expressed and associate with viral genomes between 2-3 hpi. Consistent with previous observations, ICP8 was the first replication protein to associate (Quinlan et al., 1984). By 6 hpi, replication factors are present on input viral genomes in the same relative abundance as in viral replication compartments (Figure 1C, 6 hpi + EdC), suggesting that another population of the input viral DNA is actively engaged in DNA replication during this time.

### Viral Genomes are Recognized as DNA Damage by the Host

Viral genomes enter into the nucleus as linear, naked DNA containing nicks and gaps (Wilkie, 1973). The host responds to the invading DNA by attempting to both deposit some form of chromatin on the viral genome and by triggering the recruitment of factors to initiate the repair of damaged DNA. Early during infection, the host also triggers an antiviral response, which is significantly abrogated during infection by the actions of ICP0. We visualized genomes entering into nuclei by 1 hpi (Figure 1A) and found that these genomes associate with factors that have previously been established for their role in the intrinsic response to infection. These includes components of PML NBs (PML, SP100, SUMO2), which are enriched on viral genomes by 1 hpi (Figure 2A) but no longer detected by 2 hpi (Figure 3A). By 2 hpi, PML NBs are dispersed through the actions of ICP0 (Everett et al., 1998; Everett et al., 2006). The effects PML NBs exert on viral DNA are not known. However, PML has been shown to contribute to antiviral repression in the absence of ICP0 (Alandijany et al., 2018; Everett et al., 2006).

Early during infection viral genomes also associate with factors that have known roles in the recognition or processing of DNA breaks including RPA1, PARP1, PARP14, and Ligase 3 (LIG3) (Figure 2A). RPA1 associates with input viral genomes early during infection and remains associated throughout (Figures 2A, 3A, 6A, 7A). RPA1 has been shown to be recruited to HSV-1 and human cytomegalovirus genomes during infection (Fortunato and Spector, 1998; Wilcock and Lane, 1991) and is sometimes associated with a subset of PML NBs (Dellaire and Bazett-Jones, 2004). An interesting observation from these studies is that at least 4x more RPA1 associates with input viral genomes throughout the course of infection compared to replicated viral DNA and 9x more associates with n12 input genomes compared to replicated wild type viral DNA (Table S2). It is possible that RPA1 binds to genomes that do not progress through the infectious cycle but are present in a repressed state, or that there is a unique feature of input genomes, such as nicks and gaps or ends, that enable enhanced binding of RPA1. In the future, it would be interesting to investigate the function of RPA1 during early versus late stages of infection and to determine where specifically it associates with viral DNA.

PARP1 and PARP14 add polyADP-ribose (PAR) or monoADP-ribose (MAR) groups, respectively, to target proteins. PARP proteins have been implicated in a wide variety of cellular processes including modification of chromatin, transcription regulation, DNA damage recognition and repair, and promoting inflammatory responses (Kim et al., 2005). In addition to PARP1 and PARP14, PARP9 and its binding partner DTX3L (an E3 ubiquitin ligase) were found to associate with viral genomes in the absence of ICP4 (Figure 5D). PARP9 is a catalytically inactive protein that modulates interferon gamma-STAT1 signaling (Kim et al., 2005). Further analysis of the functions of PARP proteins in viral infection is an important area for future research.

At 2 hpi, we detected IFI16 associated with input viral genomes (Figure 3) and demonstrate that detectable recruitment is dependent on the association of ICP4 with viral DNA (Figure 5). Previous studies demonstrated that the association of IFI16 with viral DNA coincides with ICP4 recruitment (Alandijany et al., 2018; Everett, 2016). IFI16 binds to HSV-1 DNA and promotes interferon β signaling (Li et al., 2012; Orzalli et al., 2012; Unterholzner et al., 2010). IFI16 is either directly or indirectly targeted by ICP0 during infection (Cuchet-Lourenco et al., 2013; Orzalli et al., 2012). However, we and others were still able to detect IFI16 associated with viral DNA in the presence of ICP0 (Li et al., 2012; Orzalli et al., 2015), suggesting that these effects are not absolute. In contrast, we did not observe recruitment to viral DNA of other factors that are known targets of ICP0 including DNA-PKcs, RNF8, and RNF168 (Lilley et al., 2010; Parkinson et al., 1999).

After the onset of viral DNA replication (3 hpi), we detected the topoisomerase TOP2A, MRN complex members MRE11 and RAD50, MMR protein MSH2, BER proteins APEX1 and XRCC1, recombination protein RECQL, and PCNA associated with viral DNA (Figure 6A and 7A). These data are consistent with our previous observations that TOP2A and MRN complex members are recruited to replicating viral DNA and that MMR proteins and PCNA are recruited to viral replication forks in a replication-dependent manner (Dembowski and DeLuca, 2015; Dembowski et al., 2017). MRE11, MSH2, and PCNA have previously been shown to be required for HSV-1 DNA replication (Lilley et al., 2005; Mohni et al., 2011 Sanders, 2015 #97). Furthermore, MRN complex members interact with the viral alkaline nuclease, UL12, and have been proposed to play a role in viral recombination during DNA replication (Balasubramanian et al., 2010).

Together these data illustrate the timing of association of specific DNA damage response and repair proteins with viral genomes. The time of association likely corresponds to the structure of the viral genome during each stage of infection. Early on the genome has ends, nicks, and gaps, which are recognized as DNA breaks by host repair proteins. At the same time, factors that mediate intrinsic responses to infection bind. These processes are countered by the actions of ICP0, which disrupts PML NBs and blocks homologous recombination by targeting RNF8 and RNF168 for proteosomal degradation. During DNA replication the genome is subject to recombination and replication-coupled repair, at which time factors that act in these processes associate. The requirement of some of these factors for productive viral infection has been demonstrated in the past. However, the functional consequences of these interactions are not known.

### A Robust Transcriptional Switch

IE proteins are expressed and subsequently bind to the viral genome by 2 hpi (Figure 1C). At this time, we observed the recruitment of several host transcription factors to viral DNA including the Mediator complex, the Integrator complex, and factors that facilitate co-transcriptional processing of RNA (Figure 3A). Mediator acts as a transcriptional coactivator of most cellular genes (Allen and Taatjes, 2015). The Integrator complex is has multiple roles in host transcription regulation including promoter proximal pause and release following initiation (Gardini et al., 2014; Stadelmayer et al., 2014), enhancer RNA biogenesis (Lai et al., 2015), and snRNA 3′ end formation (Chen and Wagner, 2010). Integrator has been shown to regulate the processing of Herpesvirus saimiri microRNAs (Cazalla et al., 2011) and to associate with nascent HSV-1 viral DNA (Dembowski and DeLuca, 2015; Dembowski et al., 2017). The observation that Integrator is enriched on HSV-1 genomes throughout infection suggests that it plays an important role in viral transcription or co-transcriptional RNA processing.

We demonstrate that robust recruitment of the Mediator complex, Pol II, transcription elongation factors, and replication proteins is ICP4 dependent (Figures 4 and 5). ICP4 is required for the expression of early viral genes, which encode the viral replication machinery, explaining the role of ICP4 in replication protein recruitment. ICP4 has been shown to interact with TFIID (Carrozza and DeLuca, 1996) and Mediator (Lester and DeLuca, 2011), and copurifies with factors involved in chromatin remodeling, transcription elongation, and RNA processing (Wagner and DeLuca, 2013). We have also shown that ICP4 is required for the binding of components of Mediator and TFIID to viral promoters (Grondin and DeLuca, 2000; Lester and DeLuca, 2011; Sampath and Deluca, 2008). Here we demonstrate that all Mediator components are either missing or significantly reduced on viral DNA in the absence of ICP4. Therefore, it is likely that the robust recruitment of Mediator by ICP4 drives Pol II recruitment and expression of early and potentially late viral genes. Taken together, ICP4 mediates a robust transcriptional switch that occurs between 1 and 2 hpi to mediate expression of early and potentially late classes of viral genes.

After the onset of DNA replication (3 hpi) another set of transcription factors are recruited, which include the Pol II kinase CDK9, TFIIH component ERCC3, and PAF complex member CTR9 (Figure 6). Recruitment of all of these factors was also dependent on ICP4 (Figure 5). It is possible that these factors play some role in promoting late gene expression after the onset of viral DNA replication or this switch may be mediated by some change in viral genome architecture that occurs at this stage of infection.

Interestingly, by 6 hpi, there is a dramatic decrease in viral genome associated transcription factors including Pol II, Mediator, and Integrator (Figure 7B). Perhaps at this time transcription is reduced on input genomes to facilitate viral DNA packaging and/or DNA replication. However, there still an abundance of RNA processing factors present. This is consistent with replication fork pulse chase data (Dembowski et al., 2017), in which transcription factors were more enriched on replication forks and RNA processing factors were more abundant on nascent viral DNA. These data may provide insight into the mechanism of replication coupled late gene transcription, suggesting that the initiation of transcription is closely linked to act of DNA replication.

Another abundant group of proteins associated with viral genomes throughout infection are factors that regulate chromatin structure, including the B-Wich, Swi/Snf, NuRD, and FACT complexes (Figure 7B). In general, the abundance of factors that regulate chromatin increase on input genomes with time. Perhaps these factors function to remove histones from replicated viral genomes or keep them from binding in the first place allowing for late gene expression. Several of the chromatin remodeling factors identified have ATPase activity including SMARCA5, SMARCA1, SMARCA4, SMARCC3, and CHD4. An additional hypothesis is that these factors act to strip proteins off of viral genomes to enable packaging of viral DNA into nascent capsids where there is a lack of proteins associated with the viral DNA.

### A Powerful Approach to Investigate Viral Infection

In this study, we present an approach to investigate viral infection from a new and powerful perspective. We define the stages in viral life cycle by the sets of viral and cellular proteins that associate with the input genome, and hence the processes that occur on it. These stages were defined from the perspective of the infecting viral genome, which because DNA replication is semi-conservative, could be tracked from when the genome first uncoats until it is packaged in progeny virions. We also utilized a virus that does not express ICP4 (n12) to identify host factors associated with a robust transcriptional switch that mediates early viral gene expression. Information regarding viral genome dynamics not only provide new insight into the involvement of viral and host proteins in processes that occur on viral DNA, but also can lead to the development of new antivirals that target these proteins.

## ACKNOWLEDGEMENTS

We acknowledge Hannah Fox and Sarah Dremel for thoughtful discussions related to this project and Frances Sivrich for technical assistance. This work was funded by NIH grants R01AI030612 and AI44812 to NAD and R21AI137652 to JAD.

## AUTHOR CONTRIBUTIONS

JAD and NAD conceived of and designed the study, NAD supervised the project and provided materials and resources, JAD developed and optimized the methodology, JAD and NAD performed experiments, JAD analyzed the data, prepared the figures, and wrote the manuscript, JAD and NAD reviewed and edited the manuscript.

## DECLARATION OF INTERESTS

The authors declare no competing interests.

## METHODS

### Cells and Viruses

Experiments were performed using MRC-5 human embryonic lung (CCL-171) or Vero African green monkey kidney (CCL-81) cells obtained from and propagated as recommended by ATCC. The viruses used in this study include the wild type HSV-1 strain, KOS, as well as the ICP4 mutant virus, n12 (DeLuca and Schaffer, 1988). n12 virus stocks were prepared and titered in the Vero-based ICP4 complementing cell line, E5.

### Virus Purification for Analysis of Virion-Associated Proteins

Confluent monolayers of Vero or E5 cells (2×10^8^ cells) were infected with KOS or n12, respectively, at a multiplicity of infection (MOI) of 5 plaque forming units (PFU)/cell. After 24 hours, infected cells were scraped into the medium. The medium containing the infected cells was adjusted to 0.5M NaCl and incubated on ice for 45 min. The cells were pelleted at 3,000xg for 15 min at 4°C. The supernatant was then filtered through a 0.8 micron filter and the filtrate was centrifuged at 25,000xg for 2h at 4°C. The virus-containing pellets were allowed to resuspend overnight in TBS. Benzonase was added to the virus sample, which was allowed to incubate for 30 min at 37°C. The virus was then layered onto a preformed 30-65% (W/V) sucrose gradient. The gradients were centrifuged in an SW41Ti rotor at 20,000 RPM overnight at 4°C. One milliliter fractions were collected from the bottom. Ten microliters of each fraction were incubated overnight in 90 μL of 0.6% SDS and 400 μg/ml proteinase K at 37°C. The digested samples were diluted 1000-fold, and 4 μl of each diluted sample was assayed for viral DNA by real-time (RT)-PCR. The peak of viral DNA corresponded to fractions just below the middle of the tube, which also corresponds to the density of HSV-1. The peak fractions were diluted with TBS and centrifuged in the SW41Ti rotor at 24,000 RPM for 2h at 4°C. The supernatant was discarded and the pellets allowed to resuspend in a small volume overnight at 4°C. Virus titer and genome number were determined by plaque assay and RT-PCR, respectively. Viral proteins were denatured in SDS sample buffer and viral protein constituents were determined by MS (Figure 1C (Virion), Figure S5, and Table S2 (Virion))

### Preparation of EdC-Labeled Virus Stocks

Confluent monolayers of 2×10^8^ Vero or E5 cells were infected with KOS or n12 virus, respectively, at an MOI of 10 PFU/cell at 37°C for 1h. After rinsing with TBS to remove unadsorbed virus, medium was replaced with Dulbecco’s Modified Eagle Medium (DMEM) containing 5% fetal bovine serum (FBS). Four hours later, EdC was added at a final concentration of 5-10 μM. Monolayers were harvested 34-36 hours after infection, freeze-thawed three times at-80°C, sonicated, and clarified by low-speed centrifugation. Viral supernatants were passed over a G-25 column to remove residual EdC. Viral titers were determined by plaque assay on Vero or E5 cells and viral genome number was determined by RT-PCR using primers specific for the viral thymidine kinase gene as described previously (Harkness et al., 2014) (Table S1).

### Viral Genome Imaging and Immunofluorescence

A total of 2×10^5^ Vero or MRC-5 cells were grown on glass coverslips in 12-well dishes. Infections were carried out using EdC-labeled virus stocks at an MOI of 10 PFU/cell in 100 μl TBS for 1h at room temperature (RT). After infection, inoculum was removed and cells were rinsed with 1 ml TBS prior to addition of 1 ml DMEM plus 5% FBS. Infections were carried out at 37°C for the indicated period of time. Cells were fixed with 3.7% formaldehyde for 15 min, washed two times with 1x PBS, permeabilized with 0.5% Triton-X 100 for 20 min, and blocked with 3% bovine serum albumin (BSA) for 30 min. EdC-labeled DNA was conjugated to Alexa Fluor 488 azide using the Click-iT EdU imaging kit according to manufacturer’s protocol (Life Technologies). Cells were rinsed with PBS plus 3% BSA, then PBS, labeled with Hoechst 33342 (1:2000 dilution) for 30 min, washed two times with PBS, then incubated with primary antibody and Alexa Fluor 594-conjugated secondary antibodies (Santa Cruz, 1:500) as described previously (Wagner and DeLuca, 2013). ICP4 antibodies include the mouse monoclonal antibody 58S (Figure 1A and S4) (Showalter et al., 1981) and the rabbit polyclonal antibody N15 (Figure S2). 58S only recognizes the dimeric form of ICP4, which binds to viral DNA (Shepard et al., 1990). Images were obtained using an Olympus Fluoview FV1000 confocal microscope. For images in Figure 1A, background subtraction and subsequent deconvolution of each Z stack was performed manually using Huygens Essential software (Scientific Volume Imaging BV). Imaris software (Bitplane AG) was used for image rendering.

### SILAC Labeling and Affinity Purification of Viral Genomes

Prior to infection, MRC-5 cells were propagated for at least three passages in medium containing heavy amino acids (L-arginine ^13^C_6_ ^15^N_4_ and L-Lysine ^13^C_6_ ^15^N_2_) while virus stocks used to infect the cells were prepared in the presence of non-isotopically labeled or light amino acids. Confluent monolayers (∽7×10^7^) of these cells were infected with EdC-labeled KOS or n12 virus at an MOI of 10 PFU/cell for one hour at RT. After adsorption, the inoculum was removed and cells were rinsed with RT TBS before SILAC growth medium was replaced. Cells were incubated at 37°C for 1-6 hours. For each sample there was a corresponding negative control, in which SILAC cells were infected with virus that was not prelabeled with EdC using the same infection conditions. Harvesting nuclei from infected cells, biotin conjugation to EdC-labeled viral genomes by click chemistry, nuclear lysis, DNA fragmentation, streptavidin purification, and elution of associated proteins were carried out as described (Dembowski and Deluca, 2017; Dembowski et al., 2017).

### Mass Spectrometry and Data Analysis

MS was carried out by MS Bioworks. The entire sample was separated ∽1.5 cm on a 10% Bis-Tris Novex mini-gel (Invitrogen) using the MES buffer system. The gel was stained with coomassie and excised into ten equally sized segments. Gel segments were processed using a robot (ProGest, DigiLab) with the following protocol. First, segments were washed with 25 mM ammonium bicarbonate followed by acetonitrile. Next, they were reduced with 10 mM dithiothreitol at 60°C followed by alkylation with 50 mM iodoacetamide at RT. Samples were then digested with trypsin (Promega) at 37°C for 4 h and quenched with formic acid. Each gel digest was analyzed by nano liquid chromatography with tandem MS (LC/MS/MS) with a Waters NanoAcquity HPLC system interfaced to a ThermoFisher Q Exactive. Peptides were loaded on a trapping column and eluted over a 75 μm analytical column at 350 nL/min, which were both packed with Luna C18 resin (Phenomenex). The mass spectrometer was operated in a data-dependent mode, with MS and MS/MS performed in the Orbitrap at 70,000 FWHM resolution and 17,500 FWHM resolution, respectively. The fifteen most abundant ions were selected for MS/MS.

For protein identification, data were searched using Mascot and Mascot DAT files were parsed into the Scaffold software for validation, filtering, and to create a non-redundant list per sample. Data were filtered at 1% protein and peptide level false discovery rate (FDR) and requiring at least two unique peptides per protein. Viral proteins with at least five spectral counts (SpC), enriched by at least two-fold over the unlabeled negative control, and present in two biological replicates were considered to be enriched on viral DNA (Table S2). In cases where no SpCs were detected in the negative control, the denominator was set to 1 to determine the fold enrichment of viral genome associated proteins. Host factors with at least 5 SpCs, enriched by at least three-fold over the unlabeled negative control, and present in two biological replicates were considered to be enriched on viral DNA. For human proteins that are common contaminants of affinity purification-MS datasets (Mellacheruvu et al., 2013), the threshold for confident detection was increased to a five-fold relative enrichment compared to the unlabeled negative control.

For SILAC analysis, data were processed through the MaxQuant software 1.5.1.0 to recalibrate MS data, filter the database at the 1% protein and peptide FDR, and to calculate SILAC heavy/light (H/L) ratios. Proteins were distinguished as either heavy or light based on a two-fold enrichment of heavy or light peptides, respectively. Proteins that fell between this range were labeled as intermediate (I). Identified proteins were displayed graphically using GraphPad Prism or Tableau software. Potential physical and functional protein-protein interactions amongst proteins identified with high confidence were illustrated using the STRING protein-protein interaction network database (Snel et al., 2000). String diagrams were modified in Adobe Illustrator for optimal data presentation.

## SUPPLEMENTAL INFORMATION

**Figure S1. Viral Protein Identification and SILAC Analysis are Highly Reproducible.** A. Viral genome associated viral proteins were detected by MS after a 1, 2, 3, or 6 hour infection with KOS-EdC or n12-EdC and SAFs of individual proteins were plotted. Proteins were distinguished as either heavy (H, red), light (L, teal), or intermediate (I, orange) by SILAC analysis. Biological replicates reveal the reproducibility of both protein identification and SILAC MS. B. Nascent capsid proteins associate with input genomes in roughly the same relative abundance as they constitute intact capsids at 6 hpi. The abundance of capsid proteins within B capsids was determined previously (Gibson and Roizman, 1972; Newcomb et al., 1993) and the graphed values correspond to the number of copies of the protein multiplied by the molecular weight. The intensity of heavy peptides was determined by SILAC analysis of capsid proteins found to associate with input viral genomes at 6 hpi. VP26 was not detected in these studies. Related to Figures 1 and 4 and Table S2.

**Figure S2. Localization of the HSV-1 Major Capsid Protein Relative to Input Viral DNA and ICP4 Throughout Infection.** Cells were fixed at indicated times after infection with KOS-EdC (1-6 hpi) or mock infection. Viral genomes (green), ICP4 (blue), and the major capsid protein VP5 (red) were imaged relative to host nuclei (dashed lines). Arrows indicate the positions of input viral DNA. The box in the corner of each image includes a 3.5x zoomed in image of the region indicated by the arrow head. Related to Figure 1.

**Figure S3. MS Analysis is Reproducible.** Comparison of the SAFs of individual proteins found to associate with viral genomes at 1, 2, 3, or 6 hpi with KOS-EdC or n12-EdC. Each point represents an individual protein with the SAF from the first biological replicate plotted on the x-axis and the SAF from the second biological replicate plotted on the y-axis. The linear regression line is shown for reference and r represents the calculated Pearson correlation coefficient, which indicates the similarity between replicate experiments. Sample KOS + EdC 6 hpi includes previously published data in which replicating viral DNA was labeled with EdC from 4-6 hpi to enable the subsequent purification of nascent viral DNA (Dembowski and Deluca, 2017). Related to Figures 2, 3, 5, 6, and 7.

**Figure S4. EdC Labeled n12 Viral Genomes can be Visualized within Infected Cell Nuclei and Form Replication Compartments When ICP4 is Supplied in Trans.** Vero or E5 cells were infected with KOS-EdC or n12-EdC and were fixed for imaging at 3 or 6 hpi. Viral genomes (green) and ICP4 (red) were imaged relative to host nuclei (blue). Related to Figures 4 and 5.

**Figure S5. KOS and n12 Virions Contain the Same Viral Protein Components.** MS analysis of viral proteins associated with purified KOS and n12 virions. Related to Figure 4. See also Table S2.

**Table S1. The Effects of EdC Labeling on Viral Genome to PFU Ratio.** Virus stocks were prepared in the presence or absence of EdC (KOS-10 μM, n12-5 μM final concentration) and genome number and PFU were determined by real-time PCR and plaque assay, respectively. n12 virus stocks were prepared and titered in the ICP4 complementing cell line, E5. Values indicate the number of genomes or PFU per μL of virus stock. Related to Figures 1 and 4.

**Tables S2. MS and SILAC data.**

## REFERENCES

Alandijany, T., Roberts, A.P.E., Conn, K.L., Loney, C., McFarlane, S., Orr, A., and Boutell, C. (2018). Distinct temporal roles for the promyelocytic leukaemia (PML) protein in the sequential regulation of intracellular host immunity to HSV-1 infection. PLoS pathogens 14, e1006769.

Allen, B.L., and Taatjes, D.J. (2015). The Mediator complex: a central integrator of transcription. Nat Rev Mol Cell Biol 16, 155–166.

Balasubramanian, N., Bai, P., Buchek, G., Korza, G., and Weller, S.K. (2010). Physical interaction between the herpes simplex virus type 1 exonuclease, UL12, and the DNA double-strand break-sensing MRN complex. Journal of virology 84, 12504–12514.

Batterson, W., and Roizman, B. (1983). Characterization of the herpes simplex virionassociated factor responsible for the induction of alpha genes. Journal of virology 46, 371– 377.

Campbell, M.E., Palfreyman, J.W., and Preston, C.M. (1984). Identification of herpes simplex virus DNA sequences which encode a trans-acting polypeptide responsible for stimulation of immediate early transcription. Journal of molecular biology 180, 1–19.

Carrozza, M.J., and DeLuca, N.A. (1996). Interaction of the viral activator protein ICP4 with TFIID through TAF250. Molecular and cellular biology 16, 3085–3093.

Cazalla, D., Xie, M., and Steitz, J.A. (2011). A primate herpesvirus uses the integrator complex to generate viral microRNAs. Molecular cell 43, 982–992.

Chen, J., and Wagner, E.J. (2010). snRNA 3’ end formation: the dawn of the Integrator complex. Biochem Soc Trans 38, 1082–1087.

Cuchet-Lourenco, D., Anderson, G., Sloan, E., Orr, A., and Everett, R.D. (2013). The viral ubiquitin ligase ICP0 is neither sufficient nor necessary for degradation of the cellular DNA sensor IFI16 during herpes simplex virus 1 infection. Journal of virology 87, 13422–13432.

Dellaire, G., and Bazett-Jones, D.P. (2004). PML nuclear bodies: dynamic sensors of DNA damage and cellular stress. BioEssays: news and reviews in molecular, cellular and developmental biology 26, 963–977.

DeLuca, N.A., McCarthy, A.M., and Schaffer, P.A. (1985). Isolation and characterization of deletion mutants of herpes simplex virus type 1 in the gene encoding immediate-early regulatory protein ICP4. Journal of virology 56, 558–570.

DeLuca, N.A., and Schaffer, P.A. (1988). Physical and functional domains of the herpes simplex virus transcriptional regulatory protein ICP4. Journal of virology 62, 732–743.

Dembowski, J.A., and DeLuca, N.A. (2015). Selective recruitment of nuclear factors to productively replicating herpes simplex virus genomes. PLoS pathogens 11, e1004939.

Dembowski, J.A., and Deluca, N.A. (2017). Purification of Viral DNA for the Identification of Associated Viral and Cellular Proteins. J Vis Exp.

Dembowski, J.A., Dremel, S.E., and DeLuca, N.A. (2017). Replication-Coupled Recruitment of Viral and Cellular Factors to Herpes Simplex Virus Type 1 Replication Forks for the Maintenance and Expression of Viral Genomes. PLoS pathogens 13, e1006166.

Dixon, R.A., and Schaffer, P.A. (1980). Fine-structure mapping and functional analysis of temperature-sensitive mutants in the gene encoding the herpes simplex virus type 1 immediate early protein VP175. Journal of virology 36, 189–203.

Everett, R.D. (2016). Dynamic Response of IFI16 and Promyelocytic Leukemia Nuclear Body Components to Herpes Simplex Virus 1 Infection. Journal of virology 90, 167–179.

Everett, R.D., Freemont, P., Saitoh, H., Dasso, M., Orr, A., Kathoria, M., and Parkinson, J. (1998). The disruption of ND10 during herpes simplex virus infection correlates with the Vmw110 and proteasome-dependent loss of several PML isoforms. Journal of virology 72, 6581–6591.

Everett, R.D., Rechter, S., Papior, P., Tavalai, N., Stamminger, T., and Orr, A. (2006). PML contributes to a cellular mechanism of repression of herpes simplex virus type 1 infection that is inactivated by ICP0. Journal of virology 80, 7995–8005.

Fortunato, E.A., and Spector, D.H. (1998). p53 and RPA are sequestered in viral replication centers in the nuclei of cells infected with human cytomegalovirus. Journal of virology 72, 2033–2039.

Fox, H.L., Dembowski, J.A., and DeLuca, N.A. (2017). A Herpesviral Immediate Early Protein Promotes Transcription Elongation of Viral Transcripts. MBio 8.

Gardini, A., Baillat, D., Cesaroni, M., Hu, D., Marinis, J.M., Wagner, E.J., Lazar, M.A., Shilatifard, A., and Shiekhattar, R. (2014). Integrator regulates transcriptional initiation and pause release following activation. Molecular cell 56, 128–139.

Gibson, W., and Roizman, B. (1972). Proteins specified by herpes simplex virus. 8. Characterization and composition of multiple capsid forms of subtypes 1 and 2. Journal of virology 10, 1044–1052.

Grondin, B., and DeLuca, N. (2000). Herpes simplex virus type 1 ICP4 promotes transcription preinitiation complex formation by enhancing the binding of TFIID to DNA. Journal of virology 74, 11504–11510.

Harkness, J.M., Kader, M., and DeLuca, N.A. (2014). Transcription of the herpes simplex virus 1 genome during productive and quiescent infection of neuronal and nonneuronal cells. Journal of virology 88, 6847–6861.

Honess, R.W., and Roizman, B. (1974). Regulation of herpesvirus macromolecular synthesis. Cascade regulation of the synthesis of three groups of viral proteins. Journal of virology 14, 8–19.

Honess, R.W., and Roizman, B. (1975). Regulation of herpesvirus macromolecular synthesis: sequential transition of polypeptide synthesis requires functional viral polypeptides. Proceedings of the National Academy of Sciences of the United States of America 72, 1276– 1280.

Ishov, A.M., and Maul, G.G. (1996). The periphery of nuclear domain 10 (ND10) as site of DNA virus deposition. J Cell Biol 134, 815–826.

Kim, M.Y., Zhang, T., and Kraus, W.L. (2005). Poly(ADP-ribosyl)ation by PARP-1: ‘PAR-laying’ NAD+ into a nuclear signal. Genes & development 19, 1951–1967.

Lai, F., Gardini, A., Zhang, A., and Shiekhattar, R. (2015). Integrator mediates the biogenesis of enhancer RNAs. Nature 525, 399–403.

Lester, J.T., and DeLuca, N.A. (2011). Herpes simplex virus 1 ICP4 forms complexes with TFIID and mediator in virus-infected cells. Journal of virology 85, 5733–5744.

Li, T., Diner, B.A., Chen, J., and Cristea, I.M. (2012). Acetylation modulates cellular distribution and DNA sensing ability of interferon-inducible protein IFI16. Proceedings of the National Academy of Sciences of the United States of America 109, 10558–10563.

Lilley, C.E., Carson, C.T., Muotri, A.R., Gage, F.H., and Weitzman, M.D. (2005). DNA repair proteins affect the lifecycle of herpes simplex virus 1. Proceedings of the National Academy of Sciences of the United States of America 102, 5844–5849.

Lilley, C.E., Chaurushiya, M.S., Boutell, C., Landry, S., Suh, J., Panier, S., Everett, R.D., Stewart, G.S., Durocher, D., and Weitzman, M.D. (2010). A viral E3 ligase targets RNF8 and RNF168 to control histone ubiquitination and DNA damage responses. The EMBO journal 29, 943–955.

Maul, G.G., Guldner, H.H., and Spivack, J.G. (1993). Modification of discrete nuclear domains induced by herpes simplex virus type 1 immediate early gene 1 product (ICP0). The Journal of general virology 74 (Pt 12), 2679–2690.

McGeoch, D.J., Rixon, F.J., and Davison, A.J. (2006). Topics in herpesvirus genomics and evolution. Virus research 117, 90–104.

Mellacheruvu, D., Wright, Z., Couzens, A.L., Lambert, J.P., St-Denis, N.A., Li, T., Miteva, Y.V., Hauri, S., Sardiu, M.E., Low, T.Y., et al. (2013). The CRAPome: a contaminant repository for affinity purification-mass spectrometry data. Nat Methods 10, 730–736.

Mohni, K.N., Mastrocola, A.S., Bai, P., Weller, S.K., and Heinen, C.D. (2011). DNA mismatch repair proteins are required for efficient herpes simplex virus 1 replication. Journal of virology 85, 12241–12253.

Newcomb, W.W., Trus, B.L., Booy, F.P., Steven, A.C., Wall, J.S., and Brown, J.C. (1993). Structure of the herpes simplex virus capsid. Molecular composition of the pentons and the triplexes. Journal of molecular biology 232, 499–511.

Orzalli, M.H., Broekema, N.M., Diner, B.A., Hancks, D.C., Elde, N.C., Cristea, I.M., and Knipe, D.M. (2015). cGAS-mediated stabilization of IFI16 promotes innate signaling during herpes simplex virus infection. Proceedings of the National Academy of Sciences of the United States of America 112, E1773–1781.

Orzalli, M.H., DeLuca, N.A., and Knipe, D.M. (2012). Nuclear IFI16 induction of IRF-3 signaling during herpesviral infection and degradation of IFI16 by the viral ICP0 protein. Proceedings of the National Academy of Sciences of the United States of America 109, E3008–3017.

Parkinson, J., Lees-Miller, S.P., and Everett, R.D. (1999). Herpes simplex virus type 1 immediate-early protein vmw110 induces the proteasome-dependent degradation of the catalytic subunit of DNA-dependent protein kinase. Journal of virology 73, 650–657.

Quinlan, M.P., Chen, L.B., and Knipe, D.M. (1984). The intranuclear location of a herpes simplex virus DNA-binding protein is determined by the status of viral DNA replication. Cell 36, 857–868.

Roizman, B., and Whitley, R.J. (2013). An inquiry into the molecular basis of HSV latency and reactivation. Annu Rev Microbiol 67, 355–374.

Sampath, P., and Deluca, N.A. (2008). Binding of ICP4, TATA-binding protein, and RNA polymerase II to herpes simplex virus type 1 immediate-early, early, and late promoters in virus-infected cells. Journal of virology 82, 2339–2349.

Sanders, I., Boyer, M., and Fraser, N.W. (2015). Early nucleosome deposition on, and replication of, HSV DNA requires cell factor PCNA. Journal of neurovirology 21, 358–369.

Sekine, E., Schmidt, N., Gaboriau, D., and O’Hare, P. (2017). Spatiotemporal dynamics of HSV genome nuclear entry and compaction state transitions using bioorthogonal chemistry and super-resolution microscopy. PLoS pathogens 13, e1006721.

Shepard, A.A., Tolentino, P., and DeLuca, N.A. (1990). trans-dominant inhibition of herpes simplex virus transcriptional regulatory protein ICP4 by heterodimer formation. Journal of virology 64, 3916–3926.

Showalter, S.D., Zweig, M., and Hampar, B. (1981). Monoclonal antibodies to herpes simplex virus type 1 proteins, including the immediateearly protein ICP 4. Infect Immun 34, 684–692.

Silva, L., Cliffe, A., Chang, L., and Knipe, D.M. (2008). Role for A-type lamins in herpesviral DNA targeting and heterochromatin modulation. PLoS pathogens 4, e1000071.

Sirbu, B.M., Couch, F.B., and Cortez, D. (2012). Monitoring the spatiotemporal dynamics of proteins at replication forks and in assembled chromatin using isolation of proteins on nascent DNA. Nature protocols 7, 594–605.

Snel, B., Lehmann, G., Bork, P., and Huynen, M.A. (2000). STRING: a web-server to retrieve and display the repeatedly occurring neighbourhood of a gene. Nucleic acids research 28, 3442–3444.

Stadelmayer, B., Micas, G., Gamot, A., Martin, P., Malirat, N., Koval, S., Raffel, R., Sobhian, B., Severac, D., Rialle, S., et al. (2014). Integrator complex regulates NELF-mediated RNA polymerase II pause/release and processivity at coding genes. Nat Commun 5, 5531.

Tognon, M., Furlong, D., Conley, A.J., and Roizman, B. (1981). Molecular genetics of herpes simplex virus. V. Characterization of a mutant defective in ability to form plaques at low temperatures and in a viral fraction which prevents accumulation of coreless capsids at nuclear pores late in infection. Journal of virology 40, 870–880.

Unterholzner, L., Keating, S.E., Baran, M., Horan, K.A., Jensen, S.B., Sharma, S., Sirois, C.M., Jin, T., Latz, E., Xiao, T.S., et al. (2010). IFI16 is an innate immune sensor for intracellular DNA. Nat Immunol 11, 997–1004.

Wagner, L.M., and DeLuca, N.A. (2013). Temporal association of herpes simplex virus ICP4 with cellular complexes functioning at multiple steps in PolII transcription. PloS one 8, e78242.

Wang, I.H., Suomalainen, M., Andriasyan, V., Kilcher, S., Mercer, J., Neef, A., Luedtke, N.W., and Greber, U.F. (2013). Tracking viral genomes in host cells at single-molecule resolution. Cell Host Microbe 14, 468–480.

Watson, R.J., and Clements, J.B. (1980). A herpes simplex virus type 1 function continuously required for early and late virus RNA synthesis. Nature 285, 329–330.

Wilcock, D., and Lane, D.P. (1991). Localization of p53, retinoblastoma and host replication proteins at sites of viral replication in herpes-infected cells. Nature 349, 429–431.

Wilkie, N.M. (1973). The synthesis and substructure of herpesvirus DNA: the distribution of alkali-labile single strand interruptions in HSV-1 DNA. The Journal of general virology 21, 453–467.

